# FLAME: Macroscopic imaging with microscopic resolution. Optical biopsy of human skin

**DOI:** 10.1101/2020.01.31.927590

**Authors:** Alexander Fast, Akarsh Lal, Amanda F. Durkin, Christopher B. Zachary, Anand K. Ganesan, Mihaela Balu

**Affiliations:** Beckman Laser Institute and Medical Clinic, University of California, Irvine, 1002 Health Sciences Rd., Irvine, California, 92612; Department of Dermatology, University of California, Irvine, 1 Medical Plaza Dr., Irvine, California, 92697

## Abstract

We introduce a compact, fast large area multiphoton exoscope (FLAME) system with enhanced molecular contrast for macroscopic imaging of human skin with microscopic resolution. A versatile imaging platform with multiple modes of operation for comprehensive analysis of live or resected thick human skin tissue, it produces 3D images that encompass sub-mm^2^ to cm^2^ scale areas of tissue within minutes. The FLAME imaging platform, which expands on a design recently introduced by our group, features deep learning, additional scanning hardware elements and time-resolved single photon counting detection to uniquely allow fast discrimination and 3D virtual staining of melanin. We demonstrate its performance and utility by fast *ex vivo* and *in vivo* imaging of human skin. With the ability to provide rapid access to depth resolved images of skin over cm^2^ area and to generate 3D distribution maps of key sub-cellular skin components such as melanocytic dendrites and melanin, FLAME represents a promising imaging tool for enhancing diagnosis accuracy, guiding therapy and understanding skin biology.

## 1. Introduction

Non-invasive clinical skin imaging with molecular contrast and high spatial resolution is important for maximizing diagnostic efficacy, guiding therapy and advancing the understanding of skin biology. Multiphoton microscopy (MPM) is a laser scanning imaging technique that has demonstrated major potential for label-free, non-invasive, skin tissue assessment due to its unique contrast, mainly based on two-photon excited fluorescence (TPEF) from NADH, FAD+, melanin, keratin and elastin fibers, and second harmonic generation (SHG) from collagen^1,2^. These fluorophores may provide additional contrast based on two-photon fluorescence lifetime detection^3,4^. In the past decade, the MPM technology has been evaluated as a clinical imaging tool in applications related to diagnosis of skin cancer^5–8^, pigmentary skin disorders^9^ and alopecia^10^, characterization and understanding of skin pigment biology^11^ and keratinocytes metabolism^12^ or monitoring the effects of cosmetic treatments^13^. While these studies demonstrate the unique clinical potential of this technology, its routine implementation into clinical practice is hindered by several technological barriers. The main challenges are related to limited scanning area and speed. The capability of imaging large areas is critical for improving diagnostic accuracy, particularly for non-uniform pigmented lesions. Melanocytes that may develop into melanoma in these lesions may be easily missed by imaging of limited areas. False-negative findings delay diagnosis and compromise treatment efficacy. Enhancing imaging speed is required in order to improve the efficiency of the imaging procedure, minimizes motion artifacts and optimizes clinical work-flow. The scientific research community employing TPEF imaging for neuroscience applications has been particularly interested in imaging large tissue areas^14,15^ at high scanning speed^16–18^. Recently, these efforts have led to the development of a two-photon mesoscope, which can produce large field-of-view images acquired relatively fast at sub-cellular resolution^19^. Its sophisticated optical design has been tailored to make this microscope ideal for brain imaging of genetically modified mouse models. However, detecting weak endogenous fluorescence signals such as those from skin fluorophores would require a significant decrease in scanning rate. Also, its large footprint is far from ideal for clinical skin imaging applications.

Successful clinical translation of the MPM technology for *in vivo* skin imaging applications requires optimization of the microscope complexity, cost and footprint in addition to significant improvement of scanning area, speed and contrast, without compromising spatial resolution. We have recently demonstrated a bench-top prototype of an MPM imaging system based on TPEF and SHG signals detection, optimized to image tissue areas of ~0.8 × 0.8 mm^2^ at speeds of less than 2 seconds per 1 MPx frame for high SNR, with lateral and axial resolutions of 0.5 μm and 3.3 μm, respectively^20,21^. In this manuscript, we present significant advances of this imaging platform, based on combining optical and mechanical scanning mechanisms with image restoration neural network computational approaches^22,23^ to allow *millimeter-to-centimeter scale* imaging within minutes, while maintaining sub-micron resolution. This imaging system also features time-resolved single photon counting (SPC) detection with sufficient temporal resolution to distinguish fluorophores, such as melanin, which facilitates its quantitative assessment, important in the diagnosis and the treatment evaluation of many skin conditions. A key component of this imaging platform is the light source, a femtosecond fiber laser, which has a small footprint that allows its integration into the imaging head. We demonstrate the performance of this compact, fast large area multiphoton exoscope (FLAME) system by *ex vivo* and *in vivo* imaging of human skin.

## 2. Methods

### 2.1 MPM imaging platform

The layout of the MPM imaging platform used in this study is depicted in Figure 1. The system is based on a customized optical and opto-mechanical design described in detail in a previous publication by our group^20^. Briefly, the optical layout includes a 4 kHz resonant-galvo scanning mechanism, custom-designed relay and beam expander optics and a 25X, 1.05 NA objective lens (XLPL25XWMP, Olympus). The optical design of this system was optimized to provide sub-micron resolution images over a field-of-view of 0.8 × 0.8 mm^2^ at speeds of less than 2 seconds per 1 MPx frame for high SNR.^20^ The low image contrast caused by the limited number of detected photons due to high scanning speed was maximized by employing sensitive PMTs (Hamamatsu R9880-20 and R9880-210) and by optimizing the signal collection optics to capture the maximum amount of radiation emitted in the epi-direction.

**Fig. 1.**
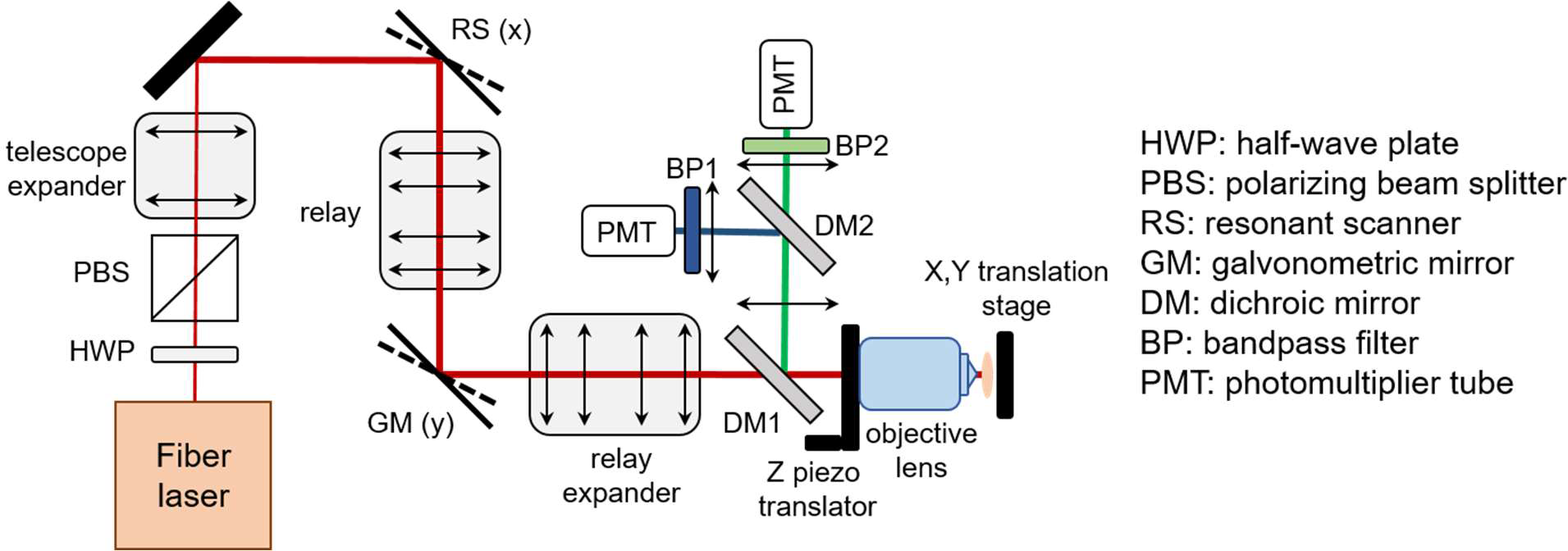
Optical layout of the MPM imaging platform scan head.

The *Results* section describes additional features we integrated in the development of this imaging platform to enhance its molecular contrast and its ability to rapidly scan up to centimeter scale areas of tissue.

### 2.2 Neural network training

We use content-aware image restoration (CARE)^22^, recently described as an effective image denoising technique, to restore fluorescence microscopy images acquired with fewer photons for enhancing imaging speed and limit light exposure time. In this method, pairs of images are acquired at low and high SNR for training a sample-specific convolutional neural network by using the high SNR images as ground truth. The trained network is then applied to restore low SNR images to predict the high SNR output. The high SNR images, serving as ground truth for training the model, were obtained by accumulating 70 consecutive frames, while the images generated by the accumulation of the first 7 frames were used as the low SNR images, the input images for the model. Our training set consisted of 128 × 128 pixel patches extracted from 900 × 900 μm^2^, 1024 × 1024 pixel images. The network was trained on 550 image pairs generated by *ex vivo* imaging of freshly excised human skin tissue (8 excisions) at different depths within viable epidermis. We used a total of 35,200 patches for the network training. Ten percent of these images (3,520) were randomly selected for the validation set, while the remaining 31,680 images were used for training. We applied this approach only to the TPEF images acquired by the FLAME system, as the SHG signal was sufficiently high at fast scanning rates. Representative input-ground truth TPEF image pair used for training the model along with the output image generated by the model are shown in Figure S1 in the Supplement 1. Training the model over 100 epochs with 30 steps per epoch resulted in a mean square error of 0.005 and a mean absolute error of 0.05.

### 2.3 Excised tissue collection and *in vivo* imaging

Skin specimens were obtained from the surgeries in the UCI Dermatology Clinic and consisted of remaining tissue from the wound closure procedures (“dog ears”). The tissue collection procedure was exempt from the Institutional Review Board (IRB) approval since the specimens were de-identified, but they were handled based on an Institutional Biosafety Committee (IBC) approved protocol. The specimens were imaged fresh, immediately upon collection. The *in-vivo* measurements were conducted according to an approved IRB protocol of the University of California–Irvine with written informed consent obtained from the subjects.

### 2.4 Data availability

The datasets generated and analyzed in the article are available from the corresponding author upon reasonable request.

## 3. Results

### 3.1 Main advances of the FLAME imaging platform

We describe below several key elements we recently introduced in the MPM imaging platform development in order to significantly enhance its performance and optimize it for clinical skin imaging. An important advance is related to the <u>excitation light source</u>. Thus, in the current design, a frequency doubled Yb-doped amplified fiber laser (Carmel X-series, Calmar Laser, USA) that generates 90 fs pulses at 80 MHz and a maximum output power of 500 mW has replaced the bulky Ti: Sapphire laser employed for excitation in our previously reported MPM platform^20^. The benefit of using this light source is that the laser head has a compact size, which facilitates its incorporation into the imaging scan head (Figure 1), allowing for enhanced compactness (35×35×20cm^3^) and portability of the exoscope.

Our recent development approaches have greatly advanced the performance of the exoscope to extend the rapid imaging with sub-micron resolution from sub-millimeter to centimeter scale. We accomplish this task by <u>employing computational approaches along with combined optical and mechanical scanning controlled by optimized data acquisition software</u>. The computational approaches, described in detail in section 3.1.1, involve the use of content-aware image restoration in order to compensate for the limited number of photons detected due to the high imaging speed. The mechanical scanning for rapid movement of the skin tissue is performed by fast, miniature linear xy stages (Q522.130, Physik Instrumente, GmbH) controlled by a customized data acquisition software (Vidrio Technologies), which synchronizes their travel with the laser beam scanning to facilitate rapid stitching of adjacent scanned areas. We selected this stage based on its performance and its reduced size (3 × 2.1 × 1 cm^3^), the latter being an essential requirement for its integration into the compact scanning head. The stage can be used as the mechanical scanner component while imaging either *ex-vivo* or *in vivo* by taking advantage of the skin elasticity. The approach of combining the mechanical and optical scanning to rapidly image sub-millimeter to centimeter scale tissue areas is described in section 3.1.2. The data acquisition software and hardware are critical components that affect the performance of the exoscope. Our current design employs the Matlab based software ScanImage (Vidrio Technologies) and data acquisition (DAQ) National Instruments (NI) hardware based on a fast digitizer (NI 5771) controlled by a FlexRIO FPGA module (NI PXIe-7975). The ScanImage software has been custom-designed for this system by Vidrio Technologies to allow the combination of optical and mechanical scanning for the multi-scale imaging described in section 3.1.2. The software enables single photon counting (SPC) detection and time-resolved SPC with a ~780 ps (16 bins) temporal resolution by using the 80MHz laser sync signal, upmultiplied 16x with a clock generator (AD9516-0, Mouser), as an external clock for the fast digitizer. Hence, detected photons may be binned into virtual output channels based on their arrival time, allowing for selective detection of some fluorophores as described in section 3.1.3, particularly of eumelanin that features significantly short fluorescence lifetime comparing to the other fluorophores in skin^4,24,25^. Although limited by modest temporal resolution, this approach provides sufficient <u>molecular contrast enhancement</u> to allow for identifying and quantifying skin pigmentation with higher accuracy.

#### 3.1.1 Enhancing imaging speed by utilizing image restoration neural network

We use content-aware image restoration (CARE)^22^ in order to compensate for the limited number of photons detected due to the high imaging speed. We trained a CARE network on pairs of TPEF images acquired at fast and slow frame rates in human skin tissue as described in the Methods section. Although the CARE network was trained on large spatial scale, low digital resolution images (0.9 × 0.9 mm^2^, 1 MPx), we found it performs just as well when applied on images acquired at high digital resolution. This is demonstrated by the TPEF images of human skin keratinocytes illustrated in Fig. 3. The 1 MPx images of 0.27 × 0.27 mm^2^ (Fig. 2a) and 0.9 × 0.9 mm^2^ (Fig. 2c) were acquired by accumulating 15 consecutive frames at an effective rate of 2 s/frame. The corresponding images predicted by the trained CARE network (Fig. 2b and Fig. 2d) show a significant enhancement of the cellular contrast, allowing for clear delineation of nuclei as shown by the line profiles (Fig. 2e and Fig. 2f) through adjacent keratinocytes in the images of Fig. 2.

**Fig. 2.**
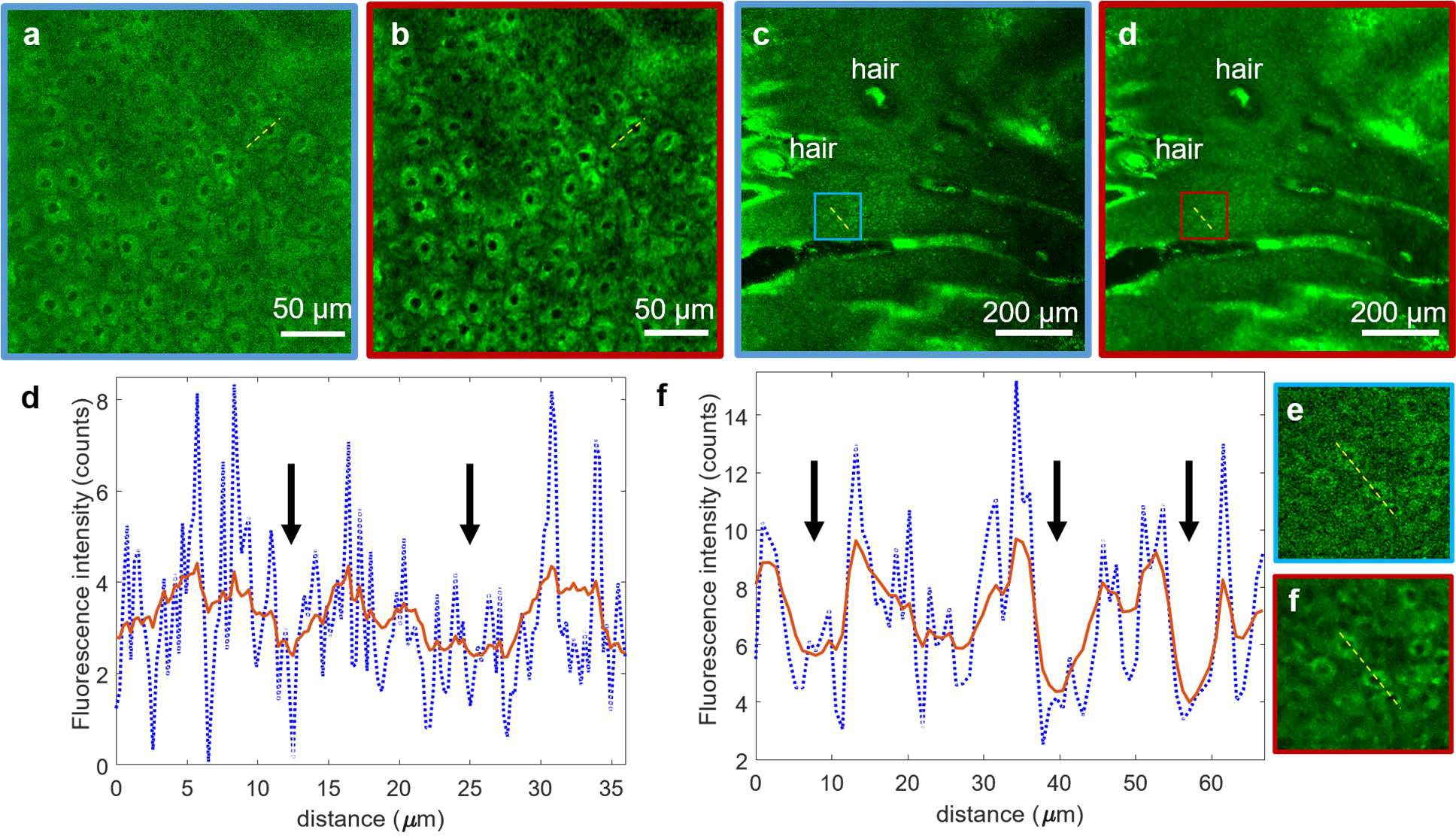
Performance of 2D CARE image restoration. Trained model is applied to images acquired by accumulating 15 consecutive frames of 0.27 × 0.27 mm^2^ (a) and 0.9 × 0.9 mm^2^ (c) image sequences (both 1 MPx, 2 s acquisition time). Restored images (b,d) display decreased noise levels and higher contrast for both fields of view. (e,f) line profiles through adjacent keratinocytes corresponding to (a,b) and (c,d) respectively (yellow lines). The line intensity profiles in the network prediction images (red) show enhanced definition of cell nuclei (black arrows) compared to the low cellular contrast to background in the raw images (blue). The insets in (g) and (h) are close-ups of the blue and red areas outlined in (c) and (d), respectively.

The content-aware image restoration has highest impact when applied to multi-millimeter scale mosaic images as it maintains/restores acceptable image contrast while using up to three-fold fewer photons. This approach provides two key benefits by reducing the acquisition time as well as the risk for potential photodamage by minimizing the exposure time.

#### 3.1.2 Laser power considerations to avoid potential photodamage

Besides minimizing exposure time through rapid scanning, we also ensure the laser fluence at the sample is below the established thermal^26^ and DNA^27^ damage threshold for two-photon microscopy of human skin. Thus, based on the focusing optics in our exoscope (NA = 1.05) and the 45 mW excitation laser power at the skin surface, the laser fluence value is lower by at least a factor of 1.7 compared to the fluence values the CE-certified clinical MPM device (MPTflex, Jenlab) employs at the skin surface and the fluence values for DNA and thermal damage threshold established for two-photon microscopy of human skin^26,27^.

#### 3.1.3 Multi-scale, high-resolution imaging by optical and mechanical scanning

For a comprehensive interrogation of the skin tissue we have devised a multiscale scanning approach that utilizes three complementary scanning modalities. We perform imaging on millimeters to centimeter scale by employing successively the following scanning mechanisms for stitching adjacent fields of views and for acquiring volumetric images:

1. <u>*the strip mosaic scheme*</u> employs the resonant galvo mirror as the fast scanner in the x direction and a miniature linear stage as the slow scanner in the y direction. The width of the strip is defined by the laser line produced by the fast scanning mirror, while its length is determined by the stage travel range. The stage moves back in the −y direction and also laterally in the x direction, a distance equal with the width of the strip, before completing each of the subsequent strips.
2. <u>*the tile mosaic scheme*</u> employs the galvo-resonant mirrors to scan each frame (‘tile’) and the linear stages to translate the sample in the x and y directions in order to scan adjacent tiles.
3. <u>*the volumetric imaging*</u> involves acquiring z-stacks of en-face images by moving the objective in the z direction, thus scanning at different depths in the skin.

Strip mosaic provides the fastest scanning rate for single frame acquisition. However, multiple frames accumulation is often required at millimeter-scale FOV for enhanced contrast to allow accurate morphological assessment. In this case, our current data acquisition software provides a faster scanning rate for the tile mosaic configuration. Figure S2 in the Supplement 1 presents a comparison of the tile and strip mosaic scanning speed for different configurations we typically employ for the multi-scale imaging.

We use these combined optical and mechanical scanning mechanisms in order to mimic the current examination method for histological tissue sections, where the pathologist examines the entire tissue at low magnification and then zooms in to investigate certain regions of interest (ROI).

Thus, in our imaging procedure, the first step of the tissue examination involves a survey scan on the centimeter scale at about 3 different depths within the epidermis, dermo-epidermal junction (DEJ) and papillary dermis. We employ the single frame <u>strip mosaic scanning mechanism</u> at this spatial scale as it provides significantly faster acquisition rate than the tile mosaic scheme (Fig. S2). We commonly acquire 10 mm long, 0.75 mm wide strips, resulting in an aspect ratio of 1:13. Depending on the sample size, we acquire up to 16 strips resulting in a scanned area of 10×12 mm^2^, typically collected as 64 MPx image and rebinned using bicubic interpolation to maintain the correct aspect ratio. The acquisition and restoration time for this type of image is 2 minutes 15 seconds. The length and the number of the strips are limited by the stage speed, its travel range (13 mm) and its synchronization with the resonant scanner. Figure 3 presents a cm-scale MPM image before (Fig. 3a) and after skin-trained CARE network application (Fig. 3b), acquired *ex-vivo* as a single frame at a 30 μm depth within the epidermis of a human skin tissue. The dynamic range enhancement obtained from neural network restoration allows for trivial field curvature correction (Fig. 3b), not feasible for low SNR raw images (Fig. 3a). The insets represent the digital zoom-in to a 200 × 200 μm^2^ area within the large-scale MPM images, showing that image restoration enhances contrast to allow visualization of keratinocytes nuclei.

**Fig. 3.**
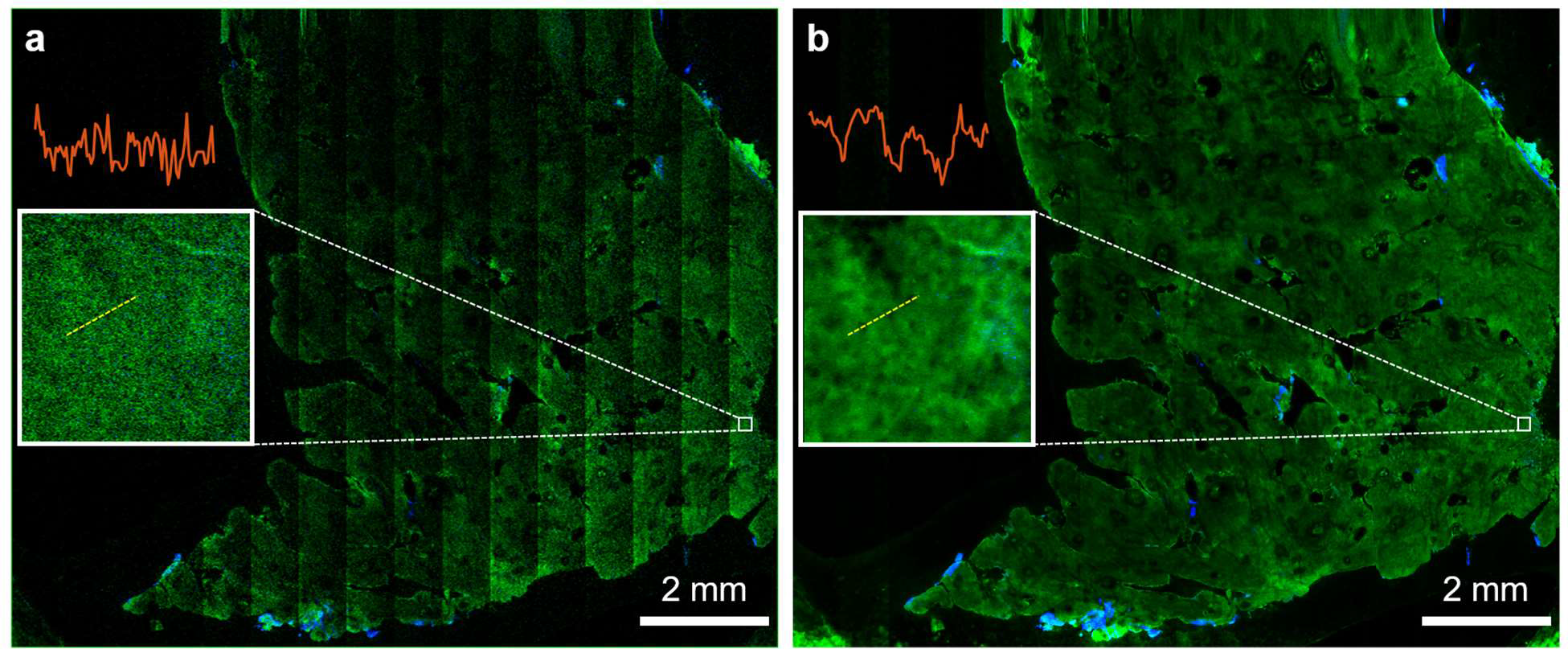
Rapid, centimeter-scale MPM *ex vivo* imaging of freshly excised facial skin at sub-micron resolution. MPM overview map (80 MPx, 1.2 × 1.0 cm^2^) of a centimeter-scale skin tissue (a) prior and (b) post CARE processing, acquired as a single frame by strip mosaic scanning at 30 μm depth in the epidermis. The insets represent digital zoom-in of the 200 × 200 μm^2^ outlined areas in (a) and (b). The intensity profiles (red) through adjacent keratinocytes show that image restoration enhances contrast to allow visualization of keratinocyte nuclei.

The images in Fig. 3 demonstrate that, by using a strip mosaic scanning mechanism and applying a trained neural network model, large areas of skin tissue can be rapidly mapped with sufficiently high resolution and contrast to allow visualization of nuclear and cellular morphology, required to identify ROIs for certain skin conditions. However, a comprehensive, detailed morphological assessment that allows visualization of microscopic features of interest, such as melanocytic dendrites, requires enhanced resolution and contrast. We accomplish this by zooming in optically (scanning millimeter scale areas) into the ROIs identified in the cm large-scale images and by accumulating (summing-up) multiple frames of the same area. We commonly employ the 15-accumulating frames <u>tile mosaic scanning mechanism</u> at the millimeter spatial scale as it provides faster acquisition rate than the strip mosaic scheme. Thus, a 4.5 × 4.5 mm^2^ area can be scanned by stitching 5 × 5 tiles of 0.9 × 0.9 mm^2^ and 1 MPx each, in 2 minutes, which includes the acquisition, the CARE network processing and stitching/display times. The high resolution and contrast of these mosaic images allow for digitally zooming in to identify architectural features and ROI selection for further volumetric scanning. A movie included in the Supplement 1 (Visualization 1) illustrates the process of navigating a high resolution tile mosaic with digital zoom and lateral browsing highlighting micron-sized melanocytic dendrites that are difficult to visualize in the fully zoomed out images.

Figure 4 illustrates the high-resolution, rapid complete mapping of a centimeter to sub-millimeter scale large human skin tissue by combining optical scanning and the two mechanical scanning mechanisms described above, followed by applying the trained CARE neural network.

**Fig. 4.**
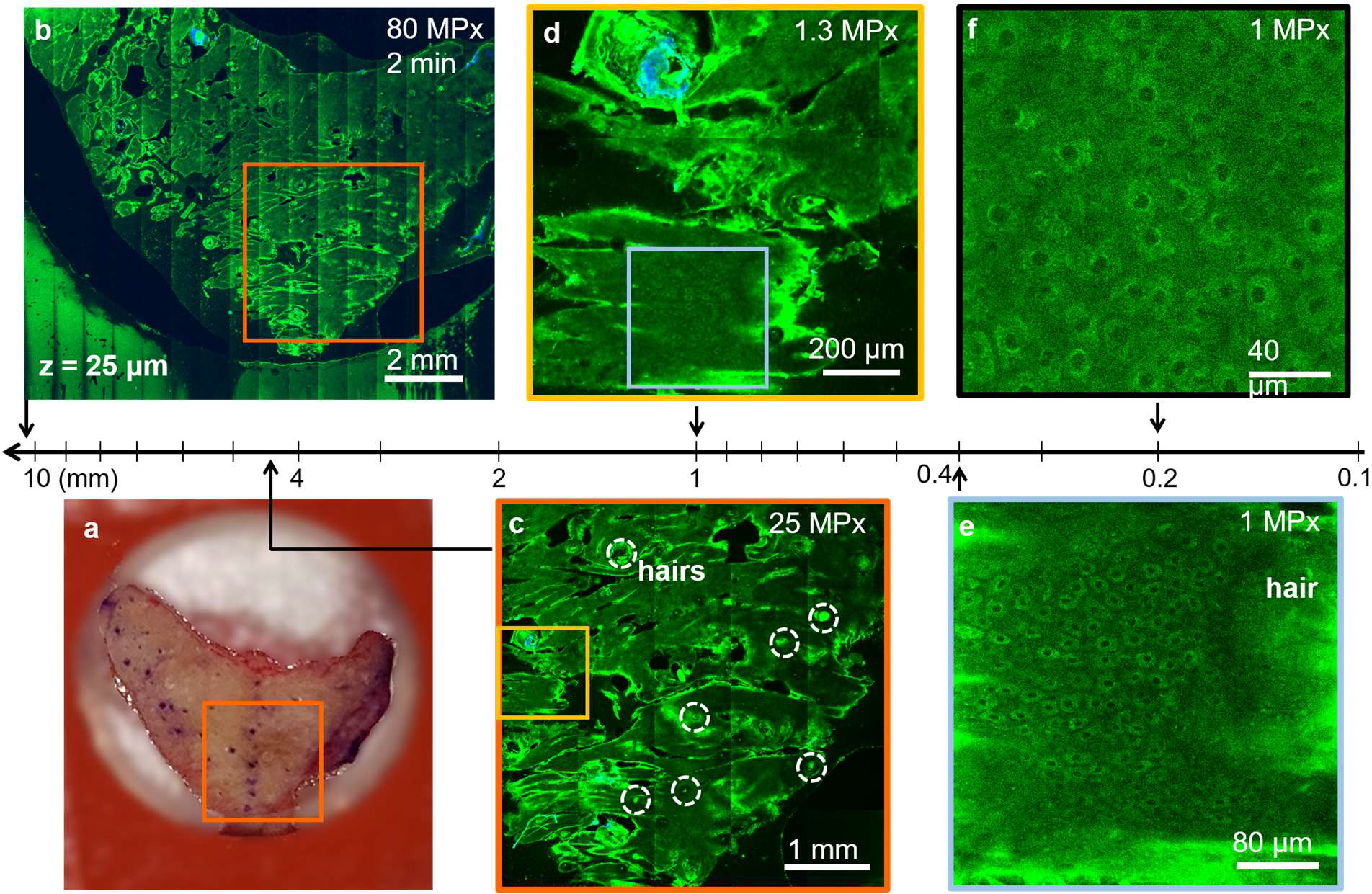
Rapid, multiscale MPM *ex vivo* imaging of freshly excised facial skin at sub-micron resolution. (a) Photographic image of the excised facial skin. (b) MPM map overview of the skin tissue epidermis generated by strip-mosaic scanning; the 80 MPx image was acquired as a single frame over a 1.2 × 1.0 cm^2^ area and restored with CARE network in 2.15 minutes. (c) Tile mosaic image of the orange outlined area in A and B covering 4.5 × 4.5 mm^2^ (25 MPx, 2 minutes) with hairs outlined by dashed circles. (d) Digital zoom in of the 1 × 1 mm^2^ yellow outlined area in C showing a close-up of the skin folds, hairs and keratinocytes within the epidermis. High-resolution (diffraction-limited) images can be obtained by “optically zooming in” i.e. by scanning sub-millimeter tissue areas with high rate sampling. (e) 1 MPx image acquired in 2s by optically zooming into the 400 × 400 μm^2^ blue outlined area in d. The image shows well-resolved keratinocytes surrounding a hair follicle. (f) Optical zoom in at higher rate sampling 1 MPx, 200 × 200 μm^2^ image allows visualization of intracellular features and protective melanin rings around the nuclei of keratinocytes (200 × 200 μm^2^, 1 MPx image acquired in 2 s).

Volumetric imaging within selected ROIs is required for in-detail assessment of the morphological changes at different depths. Z-stacks are typically acquired as a sequence of 1 MPx en-face images beneath the skin surface, encompassing volumes of about 900 × 900 × 150 μm^2^. A z-stack of 15-accumulated frames for each image, sampling a 900 × 900 × 150 μm^2^ volume every 5 μm, is acquired in 60 seconds and when needed, restored in 20 seconds. MPM images acquired *ex vivo* as a z-stack at different depths in human skin tissue are included in Supplement 1 (Visualization 2).

#### 3.1.4 Enhancing image contrast by time-resolved single photon counting (SPC)

For time-resolved SPC imaging, the signal was detected based on the fluorescence photons arrival time relative to the excitation photons, in time bins of width: −0.4−1.2 ns (red channel) and 0.4-12 ns (green channel). The rationale for defining these time bins was for attaining selective detection of melanin. The fluorescence of eumelanin, the predominant pigment in most skin types, is characterized by short lifetime with respect to most other endogenous fluorophores in skin^4,24,25^. Based on the assignment of the time bin integration in our exoscope, melanin is predominantly detected in the magenta channel, while fluorescence signals from other fluorophores such as keratin, NADH/FAD and elastin, are mainly detected in the green channel. This approach allows us to enhance the molecular contrast of the FLAME imaging platform without compromising speed. Thus, the time-resolved SPC images and the time-integrated intensity-based images require the same time for acquisition and restoration. Figure 5 illustrates millimeter-scale, sub-micron resolution MPM images based on time-resolved SPC, acquired in a normal pigmented human skin tissue at different depths.

**Fig. 5.**
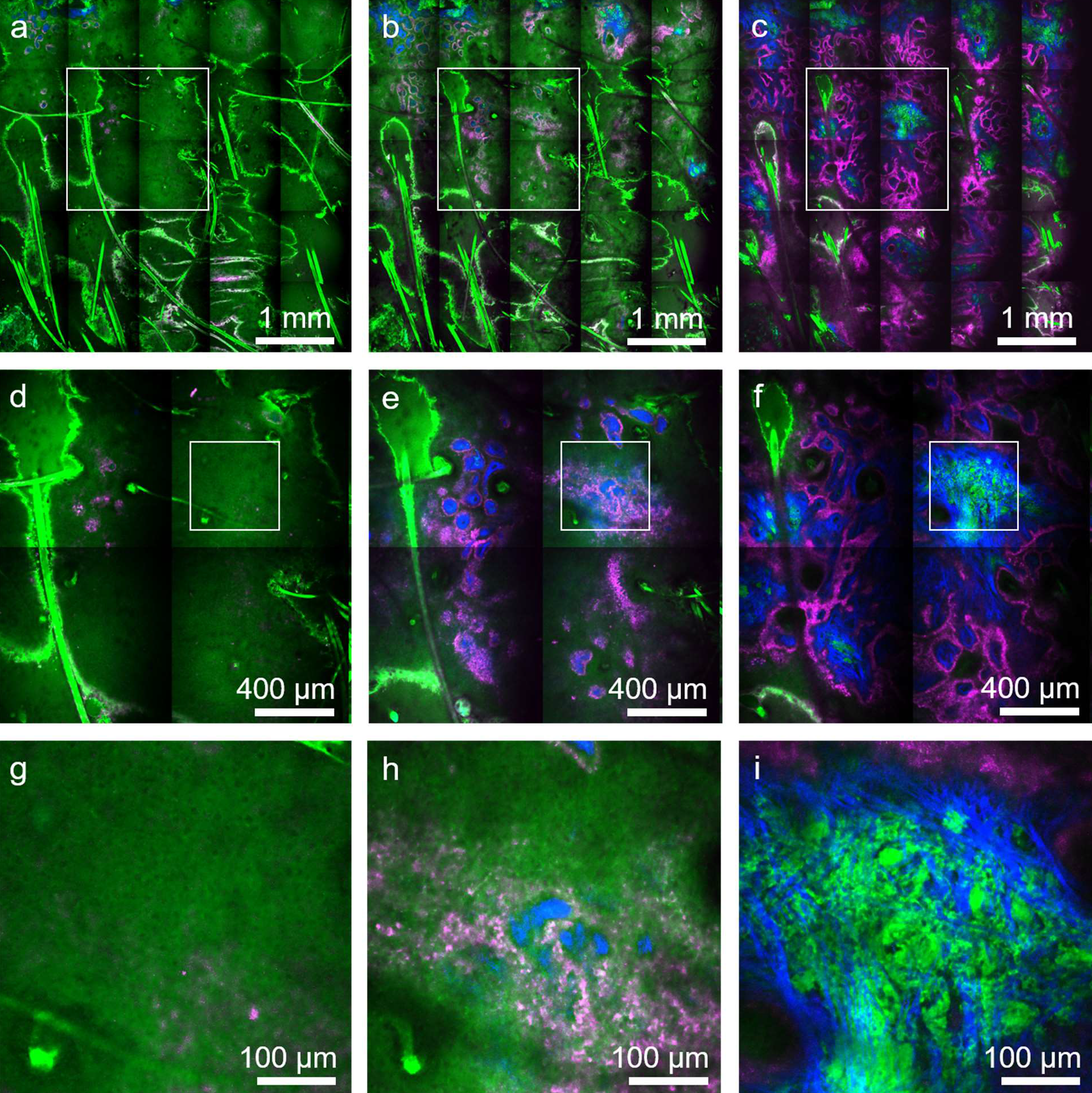
Rapid, millimeter-scale, time-resolved SPC, MPM *ex-vivo* imaging of fresh scalp tissue. Tile-mosaic images (4.5 × 4.5 mm^2^, 25 MPx) acquired in 2 minutes at select depths of 30 μm (a), 50 μm (b) and 80 μm (c) corresponding to the epidermis, DEJ and papillary dermis, respectively. Mosaic images are color-coded by short-lifetime TPEF (magenta), long-lifetime TPEF (green) and SHG (blue). (d,e,f) Digital zoom to the 1.8 × 1.8 mm^2^ white outlined areas in a, b and c, respectively, showing tissue heterogeneity on mm scale. (g,h,i) Digital zoom to the 450 × 450 μm^2^ white outlined areas in d,e and f, respectively, showing non-pigmented keratinocytes and elastin fibers predominantly visualized in the green channel, pigmented (melanin-rich) keratinocytes mainly detected in the magenta channel and collagen fibers visualized through the SHG signal in the blue channel.

Due to the integration time bin overlap of the two detection channels and the modest temporal resolution, subtraction of their intensity signals is required to obtain a differential signature of melanin. We perform the time-resolved SPC volumetric imaging using 3 channels for simultaneous detection of the SHG signal and of the TPEF signals characterized by the short and long fluorescence lifetime as described above. The 3D distribution of melanin is available almost instantaneous after correcting for signal overlap. Representative time-resolved SPC images selected from a z-stack acquired *ex vivo* in human skin are presented in Fig. 6. Figure S3 in the Supplement 1 depicts the volumetric representation that the images in Fig. 6 correspond to. The image in Supplement 1, Fig. S3 (B) illustrates the ability of the FLAME imaging system to generate 3D distribution maps of melanin, facilitating rapid quantification.

**Fig. 6.**
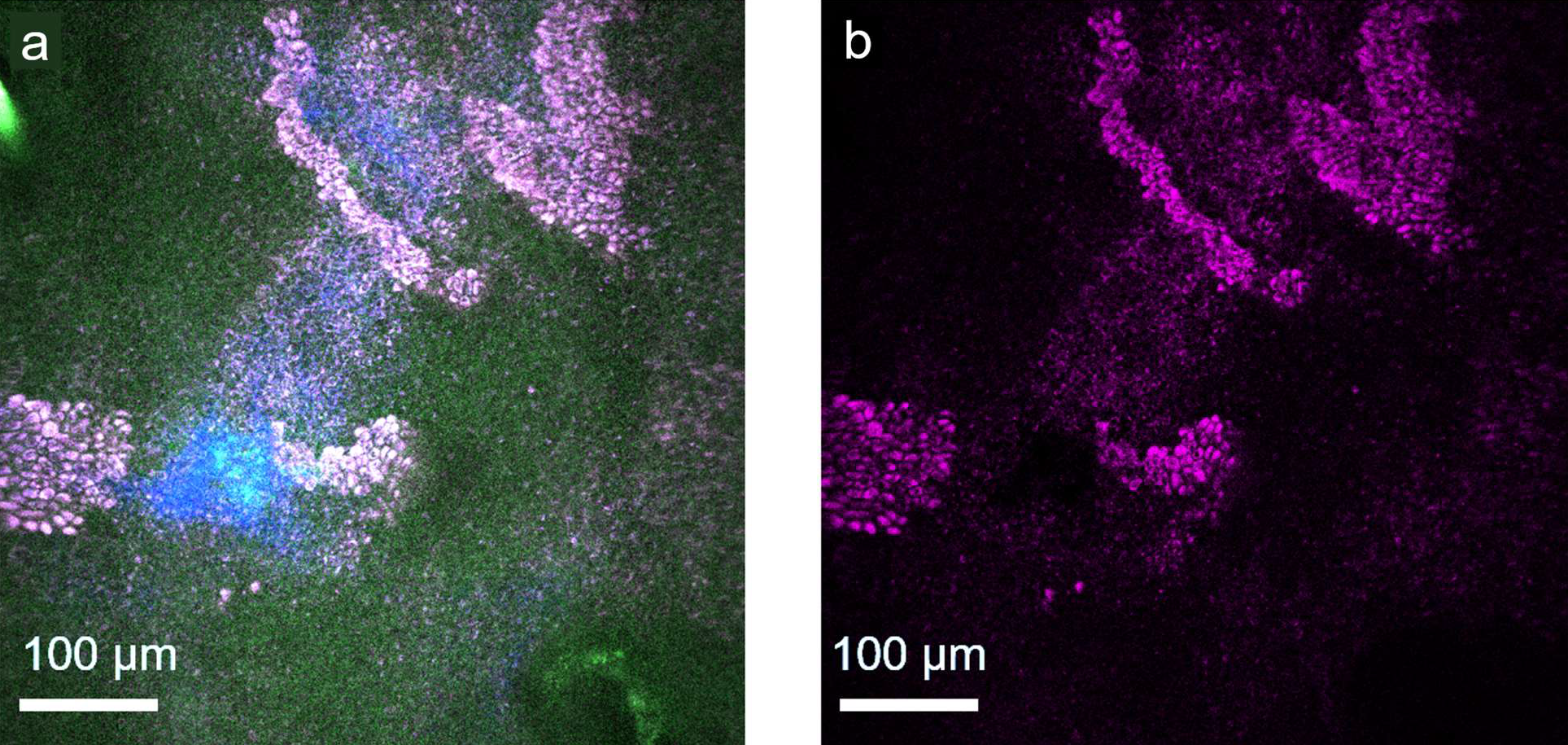
Selective detection of melanin in human skin by rapid, time-resolved SPC MPM imaging. (a) MPM image acquired *ex vivo* at 40 μm depth from the surface, at the DEJ, in a freshly excised scalp skin tissue (1 MPx, 540 × 540 μm^2^, 2 s/frame). Image shows pigmented keratinocytes (magenta-green) surrounding the tips of dermal papilla (blue). Keratinocytes are visualized by detecting their long lifetime fluorescence contributions corresponding to the green channel (mainly from NADH and keratin) and short lifetime fluorescence contributions (mainly from melanin) corresponding to the magenta channel. Collagen is visualized by detecting the SHG signal in the blue channel. (b) Melanin distribution map obtained by isolating the shortest lifetime contribution by offsetting the leakage of the green channel signal into the magenta channel from the image in a.

### 3.2 Rapid multi-scale *ex vivo* MPM imaging of human skin at sub-micron resolution

While rapid acquisition of MPM images at millimeter-to-centimeter spatial scale in human skin is critical for enhancing the efficacy and clinical utility of the MPM technology, it is essential to accomplish this goal while maintaining the ability to resolve fine molecular structures. Melanocytic dendrites are a relevant example of fine cellular components that need to be resolved in imaging applications related to melanoma diagnosis, therapy guiding of pigmentary skin disorders or characterizing and understanding of pigment biology. We demonstrate that the FLAME imaging system has the ability to resolve and rapidly map the spatial distribution of the melanocytic dendrites in the full epidermis of a freshly excised actinically damaged facial skin tissue (Fig. 7 and Visualization 1 in Supplement 1).

**Fig. 7.**
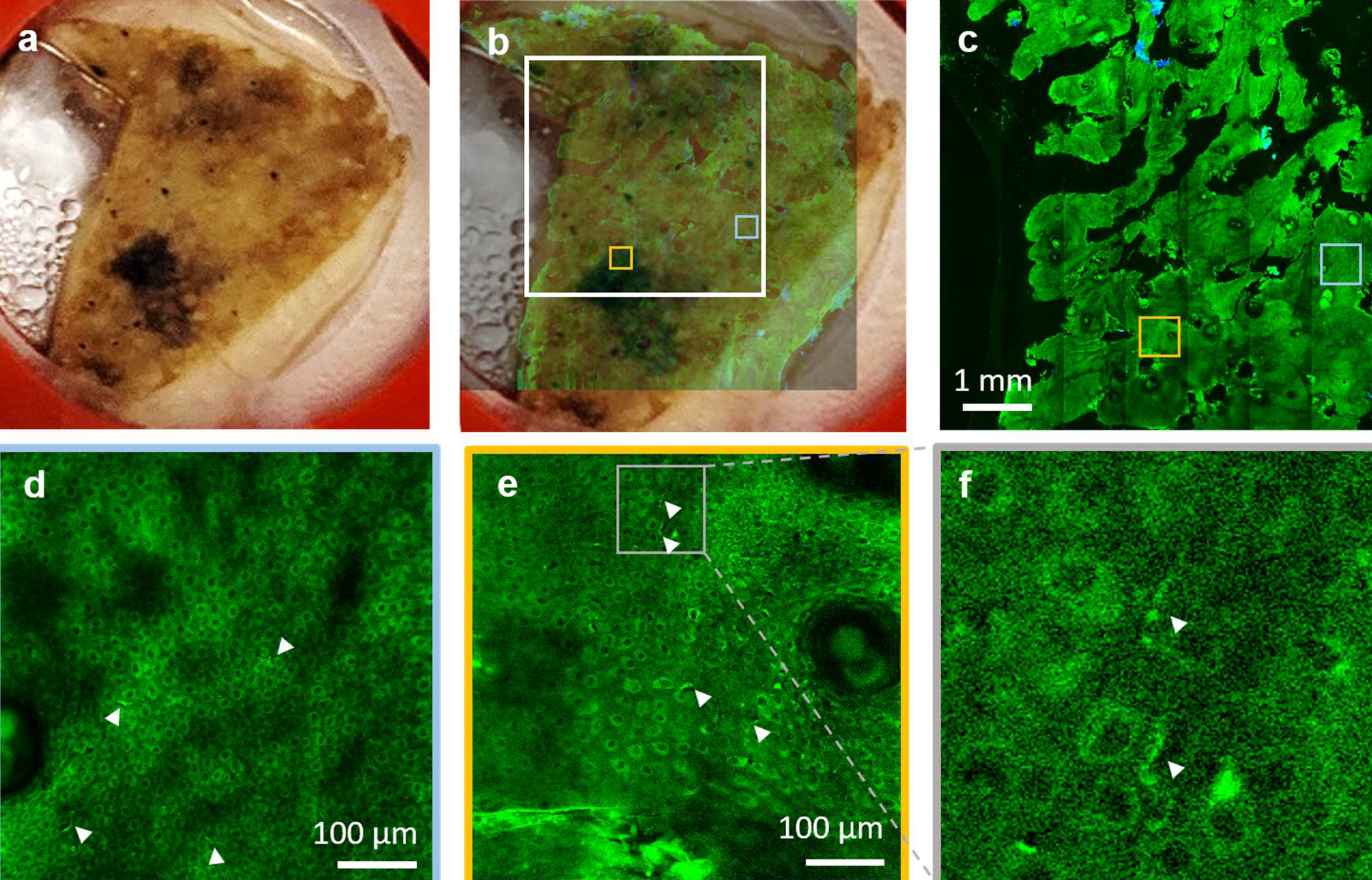
Rapid multi-scale *ex-vivo* MPM imaging with sub-micron resolution to resolve melanocytic dendrites in actinically damaged skin. (a) Photographic image of the excised facial skin tissue. (b) MPM overview map of the skin tissue epidermis generated by strip-mosaic scanning; the 80 MPx image, acquired as a single frame over a 1.2 × 1.0 cm^2^ area and restored with CARE network in 2.15 minutes, is illustrated as overlapped with the photographic image of the tissue (opacity 30%). (c) High resolution tile mosaic (6.3 × 6.3 mm^2^, 49 MPx) acquired in 3 minutes with 15 accumulated frame for each tile, at 15 μm below the skin surface. (d,e) Digital zoom into the blue and yellow outlined locations in (c), respectively. Images show the distribution of melanocytic dendrites (arrow heads). (f) A close-up of the melanocytic dendrites (white heads) in the outlined area in e.

### 3.3 Rapid millimeter-scale *in-vivo* MPM imaging of human skin at sub-micron resolution

The ultimate goal in our effort to develop the FLAME imaging platform is to utilize it as an effective imaging tool for *in vivo* skin imaging in research and clinical applications that require high spatial resolution and molecular contrast. While the imaging system is a bench-top prototype at the current stage, we demonstrate as a proof-of-concept that *in vivo* imaging using the FLAME platform is within reach. Representative MPM images with time-resolved SPC contrast acquired *in vivo* at different depths in a subject’s forearm are presented in Fig. 10. The corresponding full z-stack of 540 × 540 μm^2^ en-face images, acquired from the stratum corneum to the papillary dermis with 5 μm axial sampling can be found in the Visualization 3a, Supplement 1, along with the corresponding 3D melanin distribution (Visualization 3b, Supplement 1). The time required for acquiring and restoring 30 1 MPx frames within the z-stack was 80 seconds. A representative millimeter-scale MPM image (2.7 × 2.7 mm^2^) acquired *in vivo* from a subject’s forearm in 45 seconds, using time-resolved SPC detection, is included in Supplement 1 (Fig. S4). This is a tile-mosaic image we obtained by using a mechanical component we designed and 3D-printed to connect the miniaturized stage to the subjects’ skin.

**Fig. 8.**
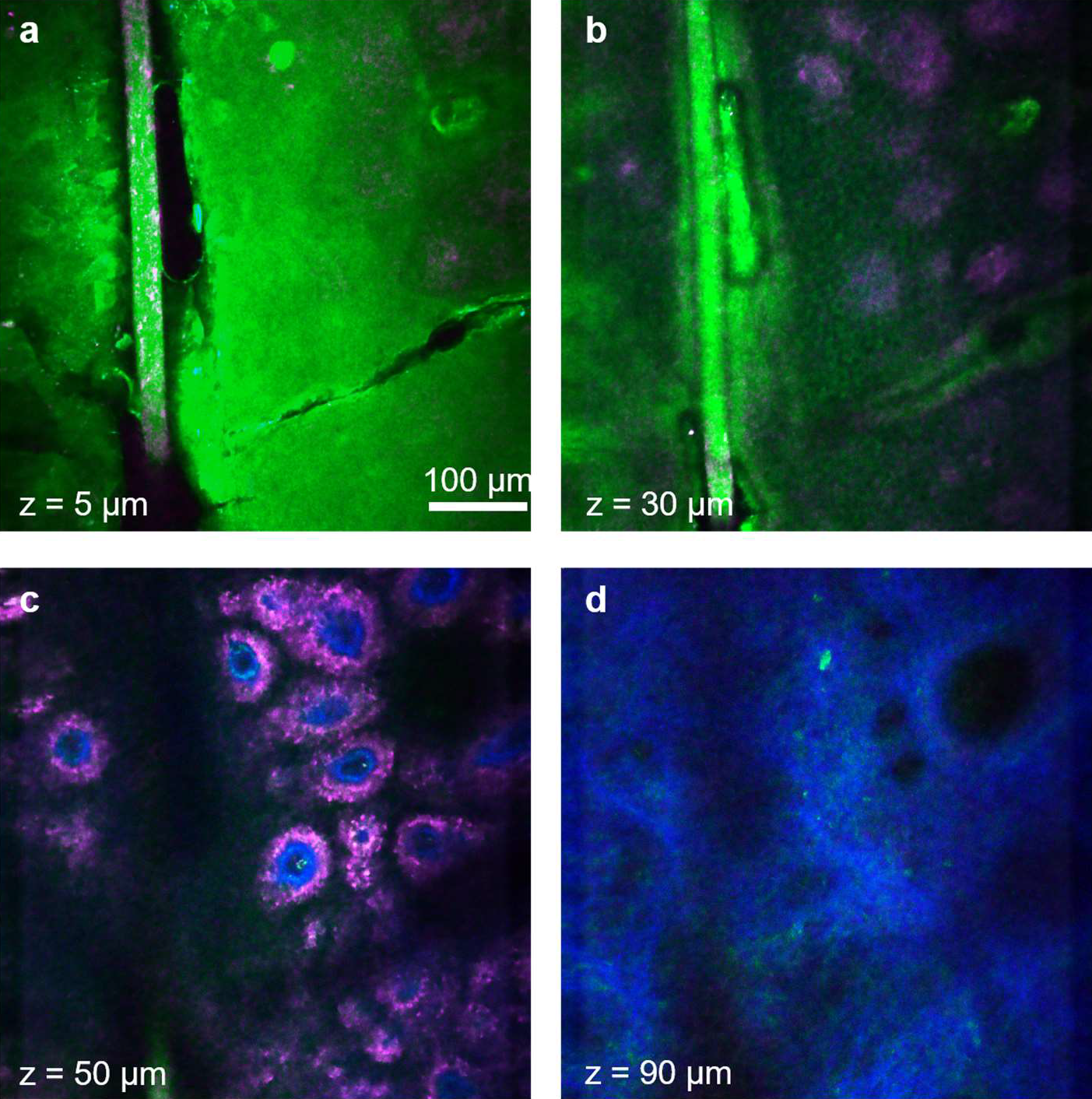
Rapid, sub-micron resolution, time-resolved SPC *in-vivo* MPM imaging of human skin forearm. MPM images acquired at different depths in human skin showing the keratinized stratum corneum (a), epidermal keratinocytes (b), pigmented cells surrounding dermal papilla (c) as well as collagen (blue) and elastin fibers (green) in (d).

## 4. Discussion

In this manuscript, we introduce a compact imaging platform based on label-free multiphoton microscopy optimized for rapid multi-scale imaging of skin tissue, providing up to sub-micron resolution, high contrast images.

One of the major challenges to overcome when performing label-free multiphoton microscopy at high scanning speed is the reduced image contrast as a result of the limited number of detected photons. We address this challenge by implementing a computational approach for image restoration in addition to our previously developed hardware optimization elements. We implemented the recently demonstrated CARE neural network approach^22^, which allowed us to enhance the image contrast and extend the scanning area to centimeter scale. Although scanning area and speed are independent parameters, they are closely related in the context of clinical translation of the MPM technology. Imaging of large tissue areas is only valuable and feasible in clinical setting when performed at high scanning rates to minimize motion artifacts and optimize clinical work-flow. We used fast, miniature linear stages and custom data acquisition software design to integrate a strip-mosaic scanning scheme. This approach, in addition to our existing tile-mosaic and volumetric imaging mechanisms, extended the spatial scanning range, allowing for rapid imaging of sub-millimeter to centimeter scale areas. A similar approach has been previously described and implemented as a solution for rapid reflectance confocal imaging, although with a much larger stage suitable for *ex vivo* imaging^28,29^.

An additional challenge to overcome in multiphoton microscopy is maintaining high spatial resolution while optimizing scanning area and speed. The high numerical aperture objective and the custom optical design of our imaging platform provide millimeter-scale images at sub-micron resolution^20^. The centimeter-scale areas are scanned rapidly at micron resolution, sufficiently high to allow, in concert with the image contrast enhancement provided by the CARE neural network model, visualization of the cellular and fibrillar structures. This performance element is critical in the ability of the exoscope to emulate the current histological examination method, where the pathologist examines the entire tissue at low magnification and then zooms in for careful investigation of selected regions of interest.

Molecular contrast is the key benefit of MPM when compared to high-resolution imaging techniques such as reflectance confocal microscopy or optical coherence tomography, currently utilized for clinical skin imaging. The molecular contrast, commonly provided in MPM skin imaging by the intensity of the endogenous fluorescence signals, can be significantly enhanced through approaches such as, fluorescence lifetime imaging (FLIM). FLIM has been demonstrated to be valuable for many applications related to skin imaging^30,31^. However, the slow acquisition speed associated with this technique represents a challenge for its potential use as a clinical tool. One solution we found for enhancing the intensity-based molecular contrast of the FLAME imaging platform without compromising speed, was to integrate time resolved SPC detection with a temporal resolution and time bins selected in such a way to allow selective discrimination of just one key endogenous fluorophore in skin, melanin. This approach is facilitated by the characteristically short fluorescence lifetime of eumelanin with respect to the other endogenous fluorophores in skin^4,24,25^. Selective detection of melanin is essential in distinguishing dermal melanocytes and melanophages from other cellular components and in applications such as pigmented lesions diagnosis and therapy assessment of pigmentary skin disorders requiring quantitative assessment of melanin.

A key element of the imaging system design is the excitation light source, a compact femtosecond fiber laser that allows its integration into the exoscope imaging head, significantly enhancing the instrument compactness and facilitating its conversion to a portable device.

We demonstrated the exoscope performance by *ex vivo* and *in vivo* imaging of human skin. The FLAME platform can provide millimeter to centimeter scale high contrast images of skin in about 2 minutes with sub-micron and micron resolution, respectively. Depth-resolved images over volumes of 900 × 900 × 150 μm^3^ are acquired in 1 minute. The spatial resolution performance was demonstrated by the ability of the exoscope to visualize melanocytic dendrites in actinically damaged human skin. Melanocytes are identified in MPM imaging by their dendritic presence. If the melanocytic dendrites are not resolved, melanocytes very often cannot be distinguished from pigmented keratinocytes. Identifying the melanocytic presence is essential in applications such as melanoma diagnosis, therapy guiding of pigmentary skin disorders or characterizing and understanding of pigment biology. An additional benefit of using FLAME as an imaging tool for these applications is related to its unique ability to provide in real-time, a volumetric mapping of melanin as demonstrated by the images we acquired *ex vivo* and *in vivo* in human skin. Rapid access to a 3D map of melanin distribution in skin is expected to expedite and enhance the accuracy of its quantitative assessment, a particularly important tool for diagnosis and therapy guiding of pigmentary skin disorders^9^ and for characterizing and understanding of pigment biology^11^.

While these features represent a significant enhancement of the scanning area and speed, molecular contrast and the overall compactness of the FLAME imaging platform, limited penetration depth in scattering tissues such as skin, still remains. Penetration depth can be optimized by using a longer excitation wavelength^32–34^, by compensating for dispersion to maintain the laser pulse duration within the tissue^35^ or by adaptive optical correction of aberrations to recover diffraction-limited performance at depth in tissue^36^. Despite this limitation, the current penetration depth of 150-200 μm is sufficient to capture early signs of malignancy in skin that occur at the DEJ^5,7,8^, to detect the presence of dermal melanocytes^7^ and to identify existing or prior inflammatory response in the papillary dermis^9,10^.

## Conclusion

We introduce a compact, multiphoton microscopy-based imaging platform, FLAME, highly optimized for rapid, label-free macroscopic imaging of skin with microscopic resolution and high contrast. It has the ability to provide 3D images encompassing sub-millimeter to centimeter scale areas of skin tissue within minutes. It allows fast discrimination and 3D virtual staining of melanin. This unique combination of features provides the FLAME imaging platform with highly optimized functionality and facilitates a seamless conversion to a portable device for rapid, multi-scale clinical skin imaging with high resolution.

## Supporting information

Visualization 1

Visualization 2

Visualization 3A

Visualization 3B

## Acknowledgements

This study was supported by the following grants: Laser Microbeam and Medical Program (LAMMP, P41-EB015890) from National Institute of Biomedical Imaging and Bioengineering (NIBIB), R01CA19546 from National Cancer Institute (NCI), R01EB026705 from NIBIB of the National Institute of Health (NIH) and by the Beckman Laser Institute programmatic support from the Arnold and Mabel Beckman Foundation. The authors also wish to acknowledge the support of the UCI Skin Biology Resource-Based Center (P30AR075047) and of the Chao Family Comprehensive Cancer Center Optical Biology Core Shared Resource, supported by the NCI of the NIH under award number P30CA062203. Alexander Fast is supported through a postdoctoral NIH Training Grant (T32CA009054). The content is solely the responsibility of the authors and does not necessarily represent the official views of the NIH.

## Author Contributions

A.F. and A.F.D designed the system components and performed the experiments. A.L. wrote the code for multidimensional data parsing, mosaic assembly and real-time image visualization. A.F. collected data, trained the CARE network and performed the image analysis. C.B.Z. provided excised de-identified tissue from surgeries and held frequent discussions regarding relevant imaging parameters for future clinical workflow integration. A.K.G. and M.B. provided guidance and advised the study. All authors reviewed and edited the manuscript.

