## Supplementary Information for "FLAME: Macroscopic imaging with microscopic resolution. Optical biopsy of human skin"

**Visualization 1: Rapid, millimeter-scale *ex vivo* MPM imaging of human facial skin with sub-micron resolution.** Movie showing a lateral browsing of a high resolution tile mosaic with digital zoom highlighting micron-sized melanocytic dendrites that cannot be visualized in the fully zoomed out images. This is a 6.3 x 6.3 $mm^2$, 49 MPx image acquired in 3 minutes.

**Visualization 2: Rapid, sub-micron resolution, *ex vivo* volumetric MPM imaging of scarred facial skin.** A z-stack of en face images at different depths covering a volume of 900 x 900 x 150 $\mu m^3$ sampled every 5 µm from the epidermis to papillary dermis. The stack of 30 1 MPx frames was acquired in 60 seconds. The stack reveals normal distribution of keratinocytes in the epidermis, pigmented keratinocytes (bright green) surrounding the dermal papilla (blue) at the DEJ and inflammatory response to scarring (outlined area) in the papillary dermis.

**Visualization 3: Rapid, sub-micron resolution, time-resolved SPC *in vivo* volumetric MPM imaging of human skin forearm.** (a) Z-stack of en face images at different depths covering a volume of 540 x 540 x 150 $\mu m^3$ sampled every 5 µm from the stratum corneum to papillary dermis. The 1MPx frames within the stack were acquired in 60 seconds. The stack reveals a bright stratum corneum with normal distribution of non-pigmented keratinocytes (green) in the epidermis and pigmented keratinocytes (magenta) surrounding the dermal papilla(blue) at the DEJ. Occasional fibroblasts (green) can be observed in the deeper layers of the papillary dermis. (b) The 3D melanin distribution corresponding to the stack shown in A, obtained by correcting for the red-green channel overlap.

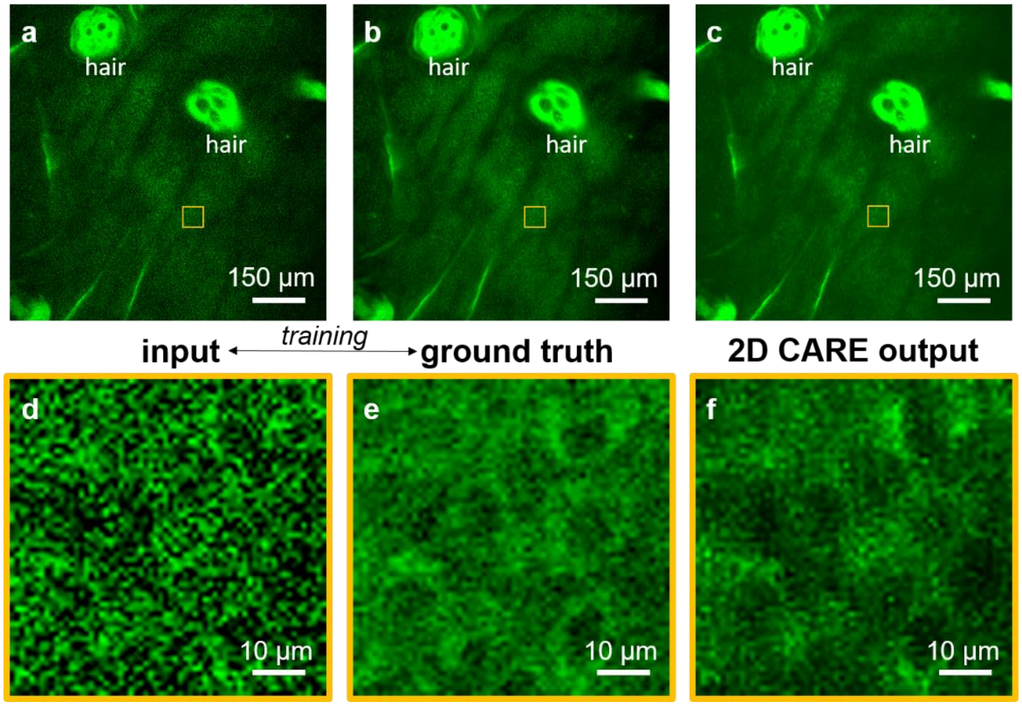


**Fig. S1. Image restoration with 2D CARE.** TPEF images (0.9 x 0.9 mm$^2$, 1 MPx) are acquired as sequential frames at 8 Hz within the epidermis of freshly excised human skin. First 7 frames of the sequence (~1 s acquisition time) are summed up and saved to serve as the network input (a). Full image sequence consisting of 70 frames (~10 s acquisition time) is accumulated to yield a single image that serves as the ground truth (b). The 2D CARE neural network output image (c) is the result of the model trained on 550 input-ground truth image pairs acquired from freshly excised human skin tissue (8 excisions) at different depths within viable epidermis. Neural network is trained on 128 x 128 pixel patches. Representative patches from (d) input, (e) ground truth and (f) network output.

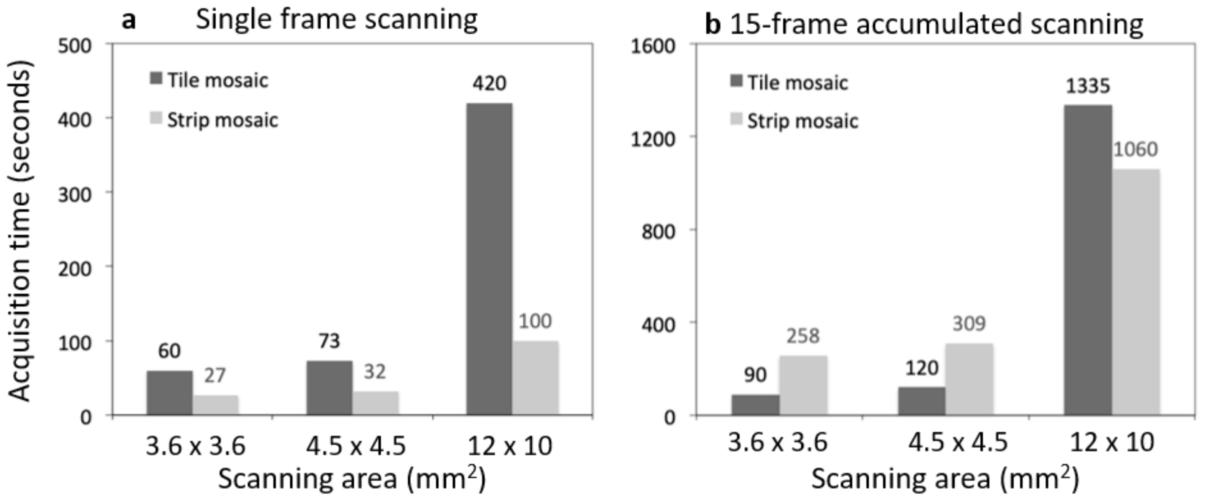


**Fig. S2. Acquisition time for different scanning areas with strip (light grey) and tile (dark grey) mosaic schemes.** Acquisition times for (a) single frame scanning and (b) 15-frame accumulated scanning corresponding to the scanning areas commonly used in our experiments.

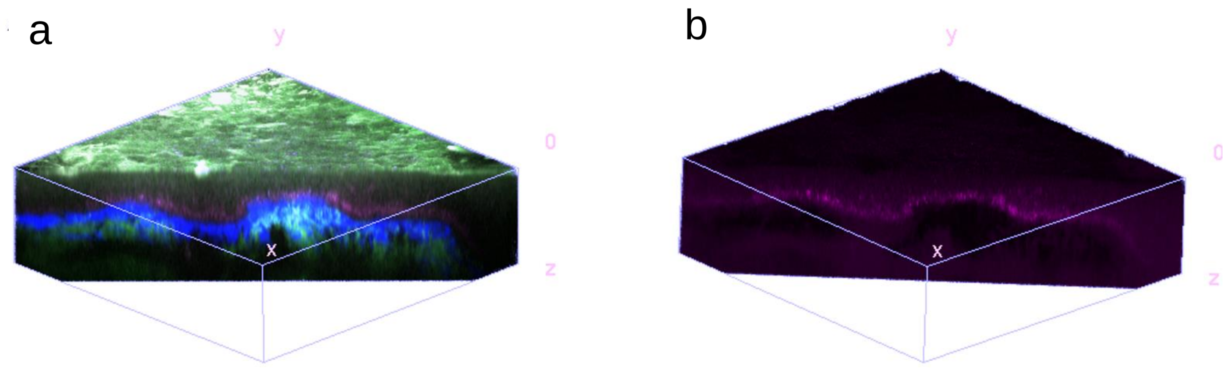


**Fig. S3. Rapid, sub-micron resolution, time-resolved SPC *ex vivo* volumetric MPM imaging of human scalp skin tissue.** Volumetric representations of a 540 x 540 x 150 μm$^3$ z-stack of 1 MPx frames, sampled every 5 μm and acquired in 60 seconds. (a) xyz-slice into the volume showing the undulations of the dermal epidermal junction. (b) 3D distribution of melanin obtained by correcting for the channel overlap (magenta channel offset by the green channel).

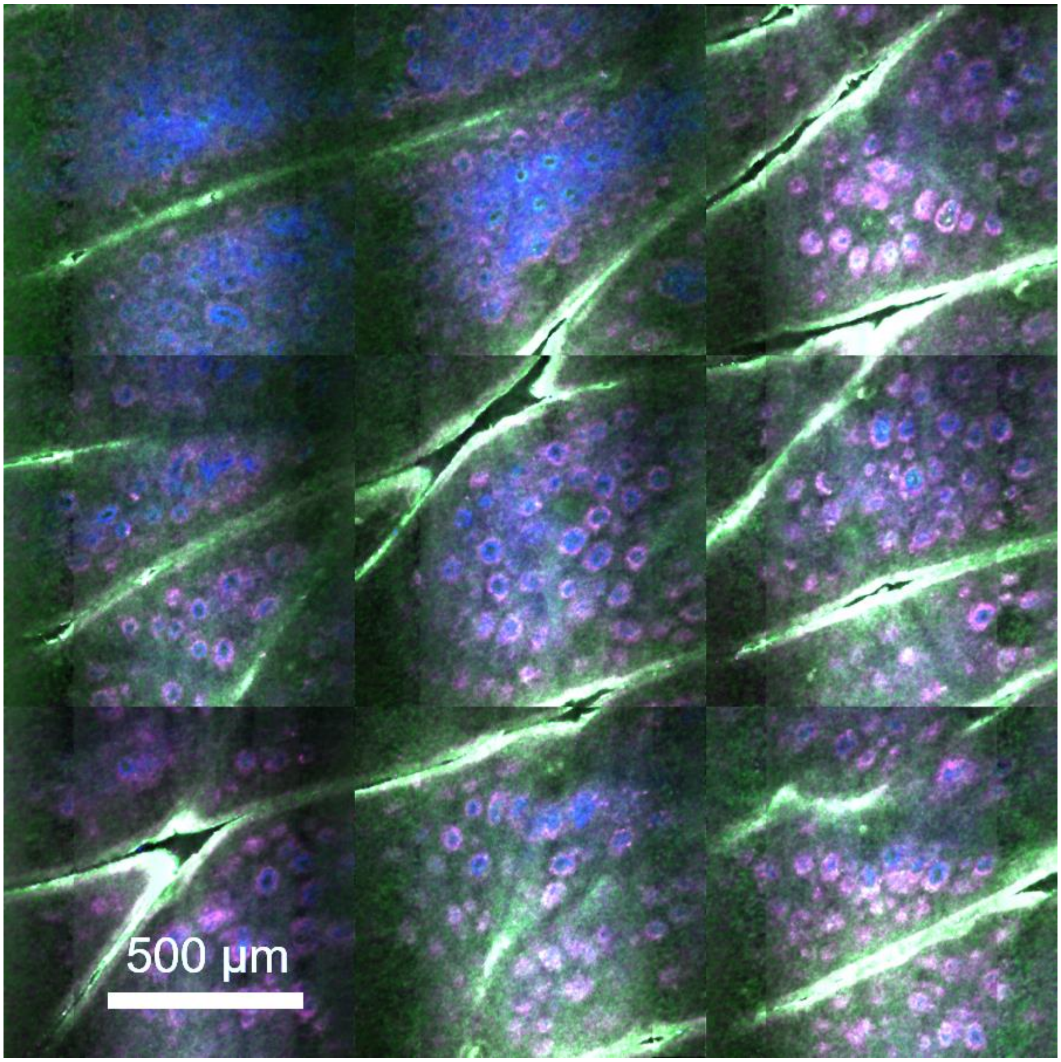


**Fig. S4. Rapid, millimeter scale, sub-micron resolution, time-resolved SPC *in vivo* MPM imaging of human skin forearm.** Tile mosaic image (2.7 x 2.7 mm$^2$, 9 MPx) acquired and restored in 45 seconds, at 45 µm below the surface at the DEJ. The image shows a map overview of normal skin morphology at this depth mainly consisting of pigmented keratinocytes (magenta) surrounding dermal papilla (blue).
